# Axonemal doublet microtubules can split into two complete singlets in human sperm flagellum tips

**DOI:** 10.1101/562777

**Authors:** Davide Zabeo, Jacob T Croft, Johanna L Höög

## Abstract

Motile flagella are crucial for human fertility and embryonic development. The distal tip of the flagellum is where growth and intra-flagellar transport are coordinated. In most, but not all, model organisms the distal tip includes a “singlet region”, where axonemal doublet microtubules terminate and only complete A-tubules extend as singlet microtubules to the tip. How a human flagellar tip is structured is unknown. Here, the flagellar tip structure of human spermatozoa was investigated by cryo-electron tomography, revealing the formation of two complete singlet microtubules from both the A-tubule and B-tubule of doublet microtubules. This different tip arrangement in human spermatozoa shows the need to investigate human flagella directly in order to understand their role in our health and disease.

## Introduction

Flagella, also called cilia, are cellular organelles that can be found in most organs in the human body as well as in many other multicellular and unicellular eukaryotes. They are membrane-covered cellular extensions that can act as a source of motility ^1–5^, a sensory organelle ^6–8^ and often a combination of both ^9–11^. Ciliopathies are a group of genetic diseases caused by malfunctions of human flagella and when such diseases specifically affect motile flagella they are called primary ciliary dyskinesia (PCD). PCD heavily affects patients’ health with recurring pulmonary infections, deafness and infertility amongst many other medical issues ^12–16^.

The flagellar proximal end emanates from a cytoplasmic basal body, which contains a ring of nine triplet microtubules (MTs) ^17–20^. Each triplet contains a complete 13-protofilament A-tubule as well as incomplete 10-protofilament B- and C-tubules. The 13-protofilament symmetry is set by the γ-tubulin ring complex in the basal body ^21–23^. At the protrusion point from the cell, the transition zone is located ^24^, where the C-tubules end while the A- and B-tubules keep extending as a doublet microtubule (dMT). In motile flagella, two MTs called the central pair (CP) start in the transition zone and extend along the dMTs, creating an ordered 9+2 MT arrangement. Together with associated protein complexes, this arrangement is called an axoneme (Figure 1A), which is the molecular machine that generates the motility of flagella. The structure of the axoneme is considered to be conserved throughout evolution ^1,4,25,26^.

**Figure 1.**
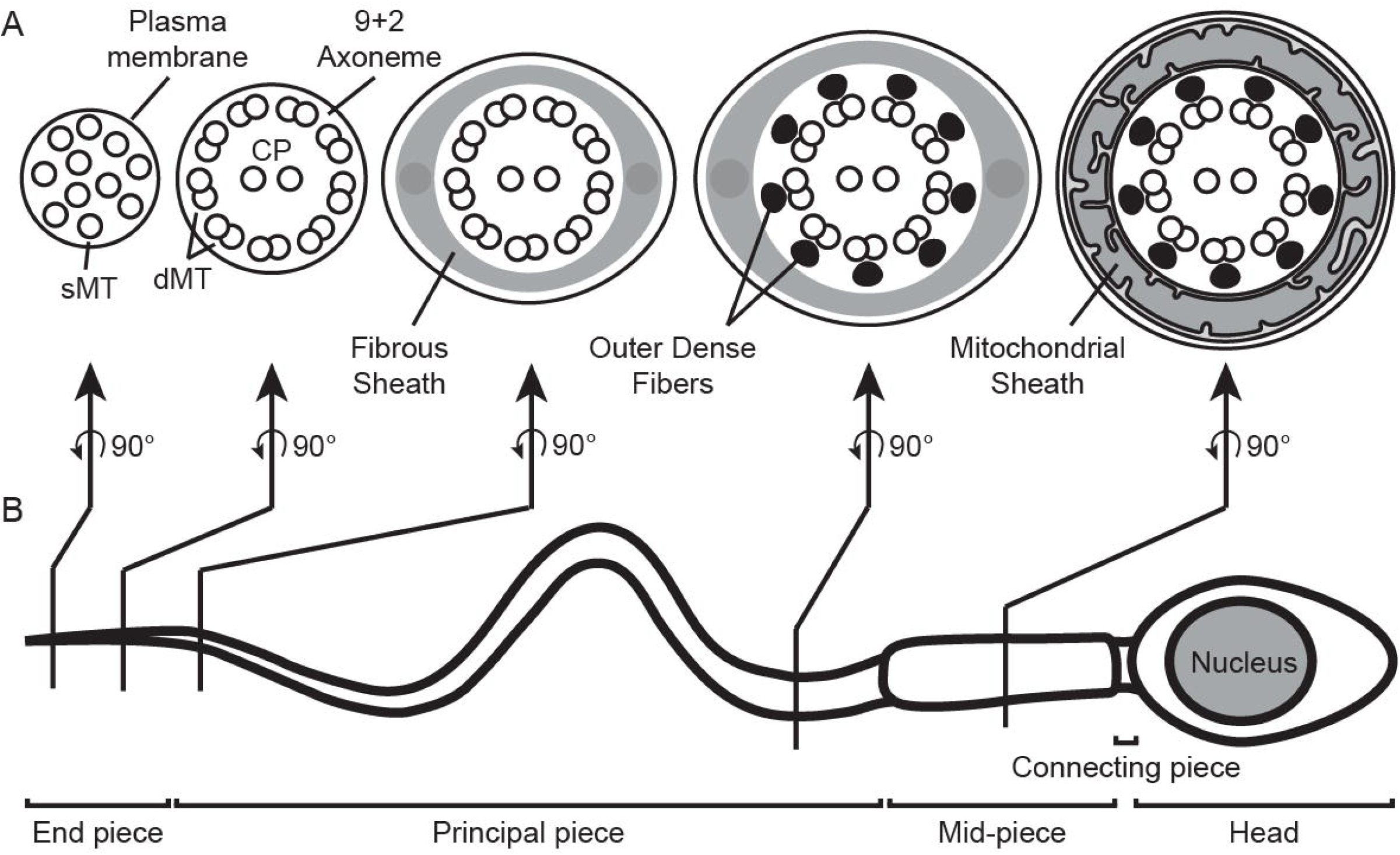
Ultrastructure of mammalian spermatozoa. (A) Cartoon cross sections through the sperm flagellum illustrate its internal ultrastructure. The sperm tail distal tip contains spatially disorganized sMTs. More proximally, MTs are organized in an axoneme with 9 dMTs and 2 CP MTs (9+2 arrangement). Even more proximal parts of the sperm tail are surrounded and protected by the fibrous sheath ^30,37^ and by the outer dense fibers, that enhance structural integrity ^33,34^ and modulate the flagellar beat ^35,36^. The mitochondrial sheath that wraps around the tail’s mid-piece provides energy for cell motility. The arrows indicate which flagellar regions that the cross sections represent. (B) The flagellum is divided into three structural regions. The most distal segment of the sperm flagellum is the end piece, which is defined by the absence of the fibrous sheath. The principal piece includes most of the tail length and is protected by the fibrous sheath. The mid-piece is defined by the presence of the mitochondrial sheath wrapped around it. The connecting piece (also called neck) links the spermatozoon head to its flagellum. The head is the most anterior part of the cell and it carries the genetic material in the nucleus.

However, a structural variability amongst flagella of different species is becoming increasingly apparent ^15,27–29^. Therefore, the information gained from model organisms may not directly apply to human flagella. Since ultrastructural information from human flagella is limited, in this study cryo-electron tomography (cryo-ET) was performed on intact human sperm tails.

In our model of choice, the mammalian spermatozoon, the flagellum is divided into three pieces (Figure 1B) ^30^. Starting at the distal tip, the end piece, the flagellum simply consists of the axoneme and the plasma membrane around it, without any supplementary structures. In the distal segment of the end piece, called the singlet region, the axonemal 9+2 symmetry is lost and the MTs extend as singlets ^13,31,32^. In the principal piece, which includes most of the flagellum length, the axonemal dMTs are supported by nine outer dense fibers, which are thought to protect the flagellum integrity ^33^, to stabilize the axonemal structure ^34^ and to modulate the power and propagation of its beat ^35,36^. In addition, the axoneme and outer dense fibers in the principal piece are surrounded and protected by the fibrous sheath, a protein matrix with a rib-like structure ^30,37^. In the mid-piece, a mitochondrial sheath wraps around the axoneme and outer dense fibers, providing energy for the motility of the cell ^30^.

The most distal tip is where the flagellum can grow and shrink. Therefore, it functions as a molecular traffic hub, where all the materials required to build the flagellum are delivered via the intra-flagellar transport (IFT) system ^38^. There is an ultrastructural heterogeneity between the distal tips of different organisms ^27,29^. In the flagellar tips of model organisms like *Chlamydomonas reinhardtii* or *Tetrahymena thermophila*, the incomplete B-tubules of dMTs end and the A-tubules extend together with the CP to the tip extremity, creating the singlet region ^39–41^. However, this region is completely absent in other models like *Leishmania mexicana* and *Trypanosoma brucei* ^29,42,43^. In rodent spermatozoa, the singlet region includes the CP together with pairs of “duplex” MTs – singlet MT (sMT) extensions of A- and B-tubules from the same dMT ^31^. The human singlet region contains up to 18 different sMTs ^13,32,36^. Since there are only 9 A-tubules and 2 CPs, any tip containing more than 11 sMTs suggests that the B-tubules in the 9+2 axoneme might be able to extend as complete sMTs as well. Our working hypothesis is therefore that the B-tubule can extend as a sMT in human sperm tails. To test this hypothesis, we examined the structure of the flagellar tips and the singlet region in human spermatozoa using cryo-electron tomography. These singlet regions revealed high variation in MT number and termination pattern between cells. Splitting of dMTs into two sMTs was directly observed, confirming our hypothesis that incomplete B-tubules can extend as complete sMTs in human flagella. Alternatives to the γ-tubulin template model to establish the conventional 13-fold symmetry of MTs have been described *in vitro* ^44,45^ and our results suggest that such alternative models may function *in vivo* as well. This study is the first to describe a human flagellar tip 3D architecture by electron tomography and the results highlight the diversity of flagellar structures across evolution.

## Results

### Variability in flagellar thickness correlates to microtubule number and presence of fibrous sheath

In order to investigate the ultrastructure of human flagella, cryo-electron tomograms were acquired on the end pieces of 23 spermatozoa. Additionally, 2D montages of cryo-electron micrographs of the sperm tail were acquired for each imaged cell, so that the position of each tomogram could be measured. A great morphological variability between different cells was observed in the montages (Figure 2A-E). Some end pieces had a constant thickness before gradually tapering to a round tip (Figure 2A), another sperm tip ended with only one sMT at the extremity (Figure 2B). Individual sperm cells also showed large variability in flagellum thickness. The thickness was measured at regular intervals from the tip in cryo-electron micrograph montages (Figure 2F; Supplementary Video 1).

**Figure 2.**
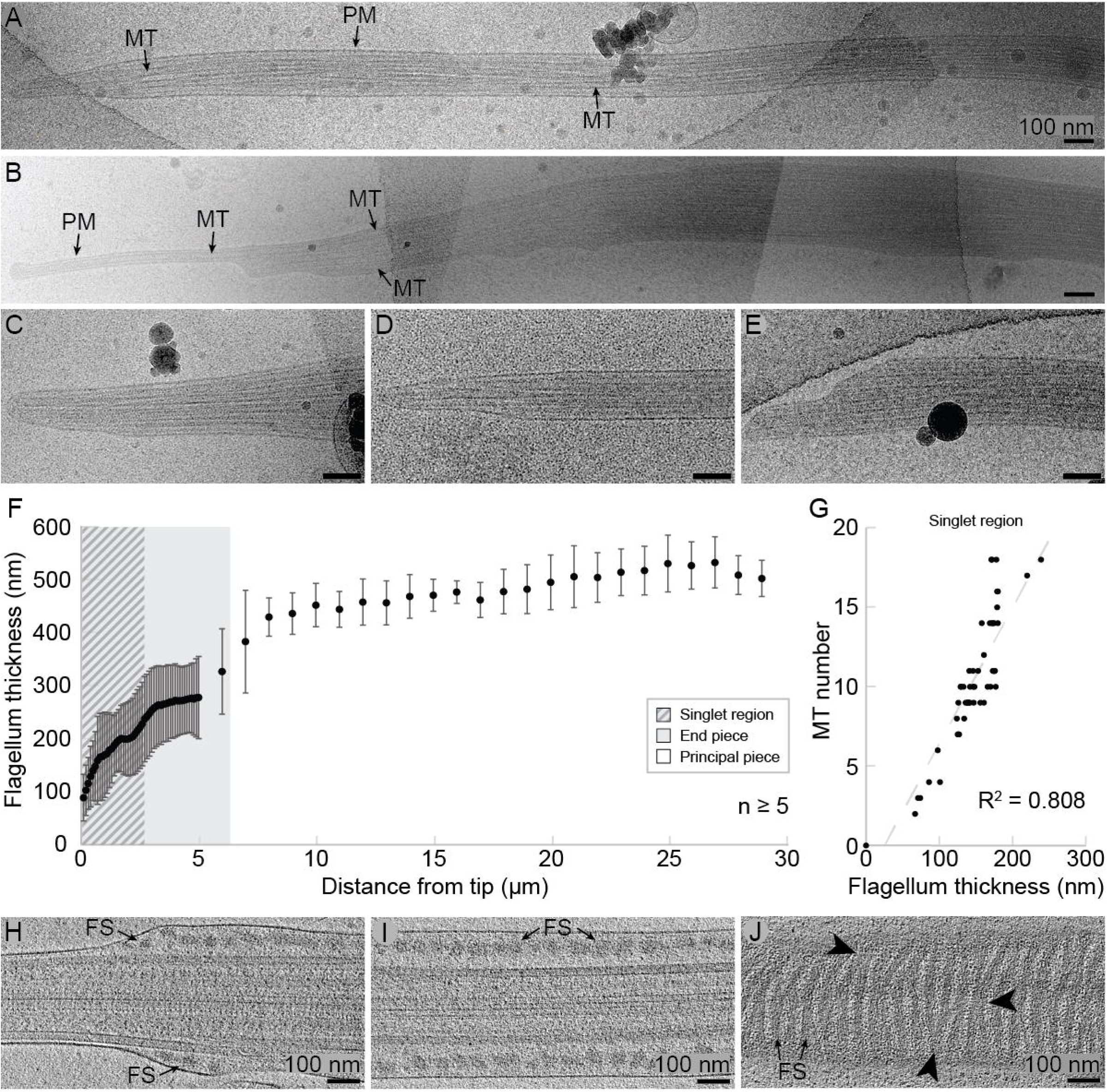
The human sperm tail thickness correlates to the microtubule number in the end piece and to the presence of fibrous sheath in the principal piece. (A-E) Montaged or single frame cryo-electron micrographs of human spermatozoon end pieces highlight morphological heterogeneity in sperm tails. The plasma membrane (PM) and the MTs of the flagellum are clearly visible. (F) The thickness of the flagellum increases with the distance from the tip. The striped background indicates the measured average extent of the singlet region and the grey background indicates the measured average extent of the end piece. Each data point represents the mean thickness of at least 5 individual flagella measured from montages of cryo-electron micrographs (up to 20 flagella per data point in the end piece). The high standard deviation (error bars) is due to the intrinsic heterogeneity of the samples. (G) Cryo-ET revealed that the MT number correlates with the flagellum thickness in the singlet region (R^2^ = 0.808). dMTs were counted as two MTs. (H-J) 23 nm-thick tomographic slices through the principal pieces of three different human sperm cells. (H) Transition point between the end piece and the principal piece. Longitudinal (H-I) and tangential (J) views of the principal piece reveal the rib-like morphology of the fibrous sheath (FS). Arrowheads indicate interconnections between adjacent ribs. All scale bars are 100 nm.

The largest change in thickness was observed within the most distal 8 µm, where the flagellum went from a tip thickness of 89 ± 44 nm (mean ± stdev) to 429 ± 36 nm (Figure 2F). In particular, the thickness mostly varied in two flagellar regions: at the distal 3 µm of the tip, including the singlet region, and approximately between 5 and 8 µm from the tip, corresponding to the transition between the end piece and the principal piece (Figure 2F). Within the principal piece, the flagellar thickness only slightly but significantly varied, reaching 501 ± 34 nm at 29 µm from the tip (*t* test between 8 and 29 µm from tip, p < 0.002). All data points had high standard deviation values, which reflects the observed high degree of heterogeneity between individual cells.

In this study, the singlet region was defined as the distance between the tip and the point where singlet and dMTs are present in equal number. Singlet and dMTs were counted for each acquired tomogram (Supplementary Figure 1) and the singlet region length was measured in three cells (2.4, 2.35 and 3.2 µm respectively), averaging to 2.7 µm. In the singlet region, no large structures other than MTs were seen and the flagellum thickness positively correlated with the MT number (R^2^ = 0.808, Figure 2G), suggesting that the MTs may be the main factor determining flagellum thickness at the tip. The MT number found in the singlet region was greatly variable, ranging from 2 to 14 sMTs, and no dMTs were ever observed closer than 1.6 µm to the tip (Supplementary Figure 1). In addition, the CP was found to terminate before other MTs in tomograms of two individual cells.

The end piece length was measured as the distance between the tip and the starting point of the fibrous sheath, corresponding to an average of 6.3 µm (n = 19). Analysis of the tomograms revealed that the increase in thickness observed at the transition point into the end piece is caused by the start of the fibrous sheath, whose structure was included in 11 tomograms at different distances from the tip (Figure 2H-J). The fibrous sheath started close to the axoneme but without any apparent auxiliary structure (Figure 2H). It then enveloped the entire principal piece with its filamentous and interconnected rib-like structure, which was visible in longitudinal (Figure 2I) as well as in tangential view (Figure 2J).

In summary, these results indicate that the number of MTs may be the main factor in determining flagellar thickness in the end piece and that auxiliary structures like the fibrous sheath are responsible for spermatozoa thickness in the principal piece.

### Doublet microtubules can split into two singlet microtubules in human sperm flagella

Three sperm tail tips were included among the acquired tomograms. In order to gain more insight in the ultrastructure of the singlet region, their MT number was counted at regular intervals from the tip, revealing diverse MT termination patterns (Figure 3A). MTs could terminate as far away from the flagellar tip as 1 µm, some extended all the way to the tip and most MTs ended approximately 300-400 nm away from it. The MT number varied between the three tips, which respectively contained 10, 11 and 14 sMTs (Figure 3A). The third flagellar tip, shown in cross-view (Figure 3B-C), stood out for having 14 complete MTs, which is more than the eleven MTs the 9 A-tubules and 2 CP MTs would form together. This observation supports the hypothesis that dMTs can generate two sMTs in human flagellum end pieces.

**Figure 3.**
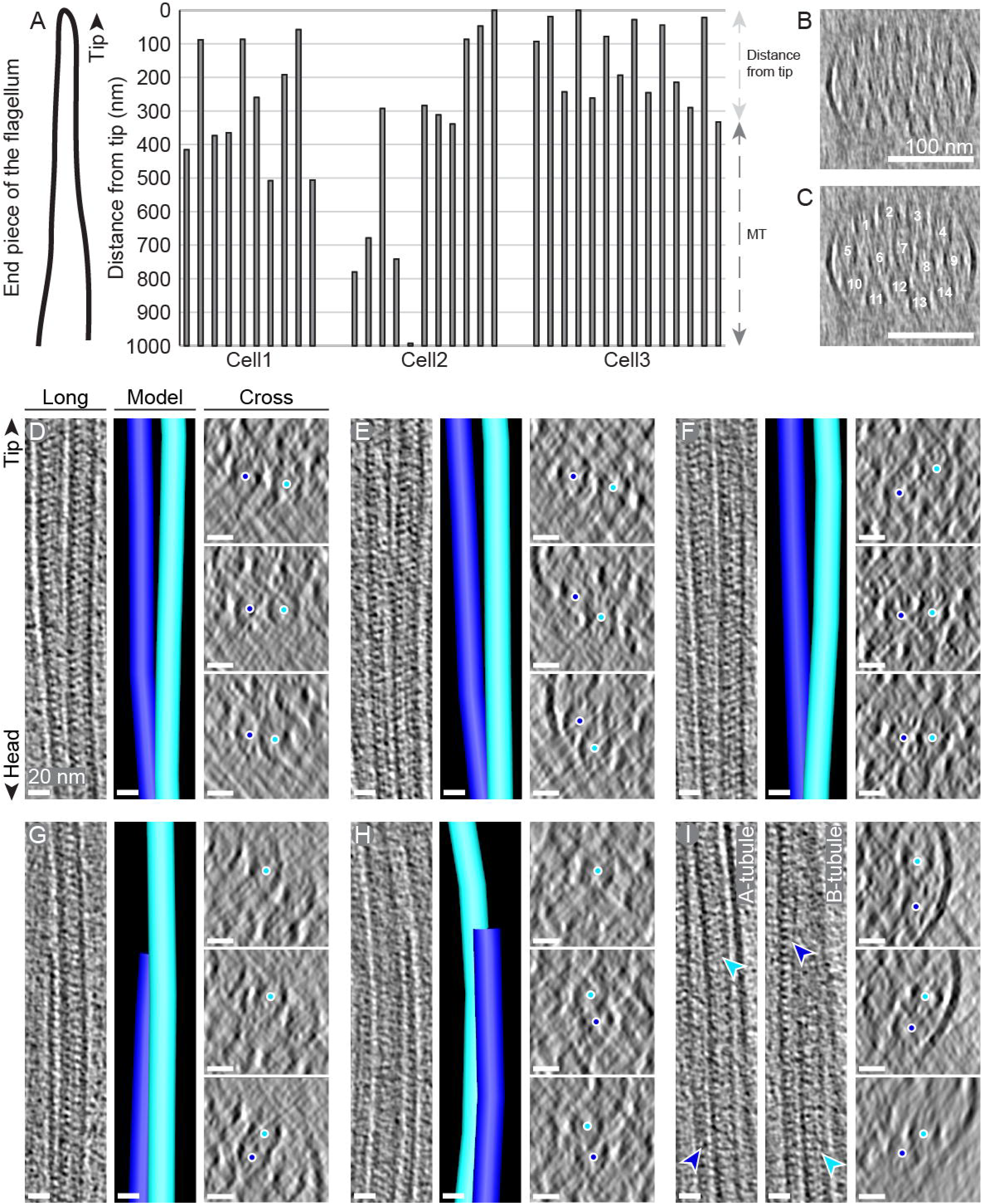
Doublet microtubules can split into two complete singlet microtubules in human flagella. (A) The distance between the MT termination point and the flagellum tip was measured in cryo-electron tomograms for each MT in the singlet region of three different sperm cells. The tips contained 10, 11 and 14 sMTs respectively. The vertical bars represent sMTs (in random order), the top of the graph represents the flagellum tip and the spacing between them represents the distance of the MT termination points from the tip. (B-C) 23 nm-thick tomographic cross-sectional slice of the singlet region in cell3 from panel A, containing 14 sMTs. The two panels show the same image, but sMTs are counted in panel C. (D-I) Doublet microtubules can transition into the singlet region by three modes. 1) The A- and B-tubules split into two complete sMT (D-F). 2) The B-tubule terminates while the A-tubule continues into the singlet region (G-H). 3) The B-tubule splits from the A-tubule and continues into the singlet region as an incomplete microtubule (I). A-tubules are drawn and marked in light blue and B-tubules in dark blue. For each dMT, longitudinal (Long) and cross-sectional (Cross) views are shown together with MT models. The cross views from top to bottom of each dMTs show areas progressively further from the tip. Panel I does not include a MT model but it shows two longitudinal views instead, one focusing on each of the splitting tubules. Longitudinal views are 8 nm-thick tomographic slices, cross views are 46 nm-thick tomographic slices. Scale bars are 100 nm in panels B-C and 20 nm in panels D-I.

In an attempt to understand which axonemal MTs the sMTs extend from, tomograms of human spermatozoa that were acquired between 1 and 3.5 µm away from the tip were examined. A tomogram acquired 2 µm from the tip revealed dMTs transitioning into the singlet region by three different modes (Figure 3D-I, Supplementary Video 2). As hypothesized, A-tubules and B-tubules originating from the same dMT split apart and continued as two complete sMTs (n = 3; Figure 3D-F). In other instances, the B-tubule simply terminated while the A-tubule extended (n = 2; Figure 3G-H). Finally, on one occasion the B-tubule split apart from the A-tubule and kept extending as an incomplete and open MT (Figure 3I). Tomograms of two other cells also included the transition of the complete axoneme into the singlet region, however they did not contain more than 11 sMTs. In these tomograms, all B-tubules terminated while the A-tubules extended to the tip, once again revealing a structural variability in the human sperm tip.

All 14 MTs in the end piece shown in Figure 3B-C were successfully used for sub-tomogram averaging in a previous study ^32^, confirming that they were all complete 13-protofilament MTs. Therefore, the *de novo* formation of three protofilaments had somehow initiated on the incomplete 10-protofilament B-tubules. In addition, while the recently discovered TAILS structure ^32^ was decorating the lumen of every MT in the tomogram with splitting dMTs, no obvious electron density was observed to specifically localize at the splitting points. We conclude that in human flagellar end pieces the B-tubule can sometimes, but not always, extend as a separate sMT.

### Doublet microtubules can also split into two singlet microtubules in bovine sperm flagella

Bovine sperm flagella were analyzed to determine whether splitting of dMTs occurs in other mammalian sperm tips as well. Electron microscopy (EM) of thin sections of high-pressure frozen bull ejaculate confirmed the presence of 18 and 15 sMTs in two different singlet regions respectively (Figure 4A-D). In addition, cryo-EM of the flattened tip of a bovine spermatozoon allowed direct visualization of dMT splitting and B-tubule termination, as observed in human flagella (Figure 4E). The analyzed tip contained at least 16 sMTs, 5 distinct dMT splitting events and one B-tubule termination event. Therefore, different modes of dMT transition into the singlet region, including extension of a B-tubule singlet, are conserved in both bovine and human spermatozoa.

**Figure 4.**
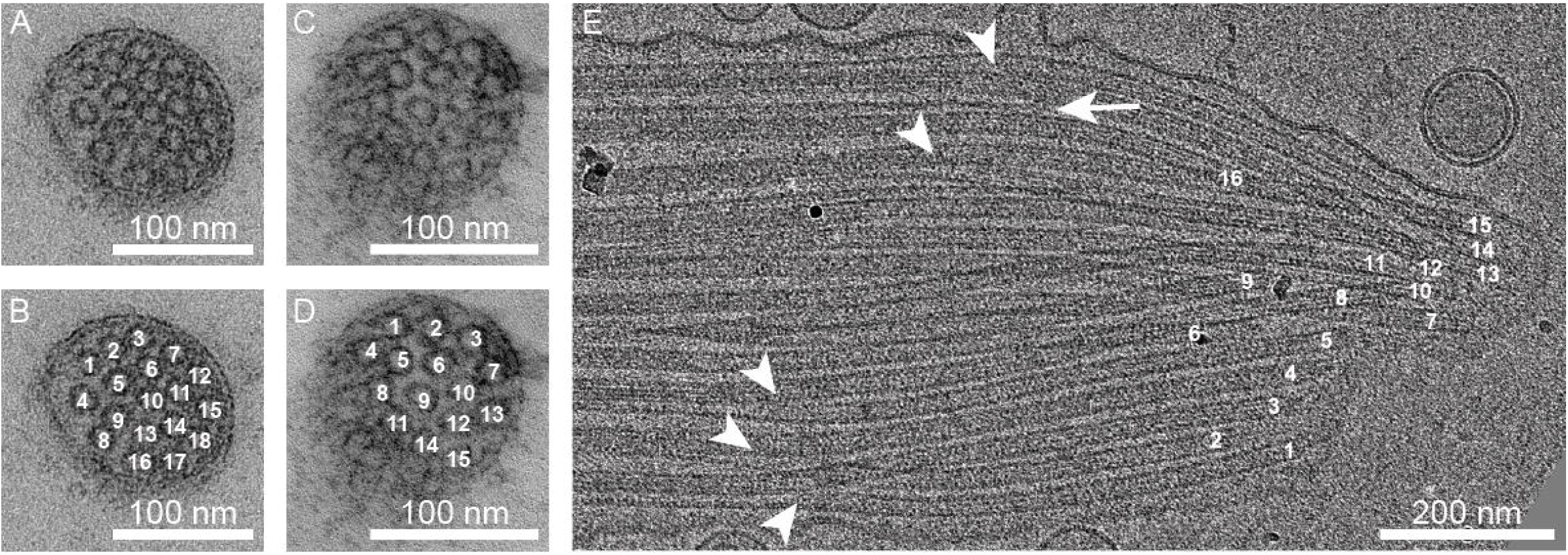
Splitting of doublet microtubules in bovine sperm flagella. Thin sections through the singlet region of bovine sperm cells show at least 18 (A-B) and 15 (C-D) sMTs. Panels A-B and C-D respectively show the same images, but sMTs are counted in panels B and D. (E) Cryo-electron micrograph of a flattened bovine sperm flagellum tip. Five events of dMTs splitting into two sMTs are shown (arrowheads) and at least 16 sMT ends are visible (numbered). One event of B-tubule termination is shown by the white arrow. Scale bars are 100 nm in panels A-D and 200 nm in panel E.

## Discussion

This study presents the first cryo-ET analysis of the cellular structure of intact human flagella, in particular human sperm tails. 3D reconstructions of close to natively preserved sperm cells revealed that their distal tips are greatly heterogeneous and not directly comparable to model organisms such as *C. reinhardtii* or *T. brucei*. For instance, in human spermatozoa the CP can terminate before any other axonemal MT (Supplementary Figure 1), unlike in *C. reinhardtii* ^40,41^, and a long singlet region is present (Figure 2A-F), unlike in *T. brucei* ^29,43^.

We observed a higher number of MTs in the singlet region of human spermatozoa than what would be expected if only the A-tubules and CP extended to the tip (Figure 3B-C) and, as it had previously been suggested ^13^, we hypothesized that dMTs could split into two complete 13-protofilament MTs. An alternative explanation for this observation could be that broken MT fragments at the flagellar tip might shift longitudinally due to the beat of the flagellum. This would lead to a higher MT count for the flagellar regions where the broken fragments are located. However, only distal MT terminations were observed in our tomograms, therefore we consider this second scenario to be unlikely. Our hypothesis that dMTs can split into two sMTs was also confirmed by the direct visualization of dMT splitting events both in human and bovine spermatozoa (Figure 3D-I and 4E). Sub-tomogram averaging of the sMTs at the sperm tip confirmed that all MTs had 13 protofilaments ^32^. Therefore, the formation of three additional protofilaments on the incomplete B-tubule is initiated *de novo* at the splitting locus. Currently unknown structures might be responsible for inducing these splitting events and for the formation the three new protofilaments. We do not know of any protein that separates dMTs, but a hypothesis is that the plus-end tracking protein EB1 helps the initiation of three new protofilaments, since it has been reported to promote 13-protofilament MT formation ^44^ and tubulin sheet closure ^46^ *in vitro*. The small size of EB1 (30 kDa) ^47,48^ would further speak for this, since we saw no distinct electron densities at the site of dMT splitting. Alternatively, the currently uncharacterized TAILS structure ^32^ may somehow be involved in these processes. An additional hypothesis could be that a larger complex only temporarily assembles during sperm maturation to split the dMTs.

What advantage might the spermatozoa gain by splitting dMTs? Since the number of splitting events does not seem to be consistent between cells (in fact some do not split at all), and since the *de novo* nucleation of protofilaments in B-tubules might not be 100% efficient (Figure 3I), it is possible that they may serve as a source of variability. A high morphological intra-individual heterogeneity in human spermatozoa has been previously reported in terms of ultrastructure ^36^, motility and viability ^49^. Given the large number of sperm cells that are produced in mammals, this may be an evolutionary strategy to ensure that some will succeed in fertilization. Another advantage of inducing sMTs might be related to the newly discovered TAILS structure, which was hypothesized to stabilize MTs in the singlet region ^32^. It is possible that TAILS may stabilize sMTs more strongly than dMTs, therefore a transition from dMTs to sMTs would maximize MT stability.

The fact that more than 11 MTs exist in the human sperm singlet region has been observed previously ^13,32,36^. In rodent sperm, dMTs and the CP were reported to form “duplex” MTs that extended to the flagellar tip, where they were tied together by capping structures ^31^. This suggests that splitting of dMTs may be a common phenomenon among mammals. However, contrary to rodent spermatozoa, no “duplex” structure was observed in human flagella. MTs in the human singlet region did not terminate in pairs (Figure 3A) and no apparent structures tied MTs together.

The CP was observed to terminate as far as 7.4 µm from the tip, in the principal piece (Supplementary Figure 1) ^32^. To the best of our knowledge, no human singlet region with more than 18 MTs (which possibly originated by splitting of all 9 dMTs) has been reported. Therefore, we speculate that the CP in human spermatozoa may generally terminate before any other axonemal MT, contrary to the model organism *C. reinhardtii*, where the CP extends the furthest into the flagellar tip ^40,41^. This would imply that the flagellar motion, which is modulated by the CP in spermatozoa ^50,51^, may differ in the end piece from the rest of the flagellum. However, due to the relatively short extent of the sperm tail included in each tomogram, it was impossible to track the CP MTs through the entire end piece of every imaged cell. Therefore, we cannot exclude that some MTs observed in tomograms of singlet regions are in fact the CP. A more extensive tomography study or immuno-electron microscopy against specific CP MT markers would address this question more appropriately.

We provide here the first detailed cellular 3D architecture of a human flagellar tip, revealing extensive flagellar structural variability compared to the commonly used model organisms. However, it remains to be investigated if the flagellar tip structure is conserved between different cell types in humans, for example by comparing to ciliary tips found in the upper respiratory tract. Revealing the human 3D flagellar structure will inform us of which model organism best resembles the human flagellum and should be prioritized in future ciliary research.

## Materials and Methods

### Sample collection and plunge freezing

Human sperm was donated by three healthy men and plunge-frozen within 1-3 hours from ejaculation time. Bull sperm was collected by VikingGenetics (Skara, Sweden), diluted in OptiXcell medium (IMV Technologies), stored at 4°C upon arrival to the lab and frozen for EM within 4 days from collection. A Vitrobot climate-controlled plunge freezer (FEI, Eindhoven, Netherlands) was used for plunge freezing of human and bull sperm. Prior to plunge-freezing, 1µl of gold fiducials was mixed in 4 µl of ejaculate and the 5 µl mix was blotted for 3 s (human) or 5 s (bovine). Leftover cells were examined under the light microscope, where their motility ensured that viable spermatozoa had been frozen.

### High-pressure freezing and freeze substitution

Bovine spermatozoa were prepared using high-pressure freezing followed by freeze substitution. Undiluted bull ejaculate was mixed with an equal amount of 1x PBS and then concentrated by centrifugation at 2000 x g for 30 seconds. The supernatant was removed and cells were washed with 1x PBS and concentrated again. Concentrated cells were loaded onto aluminum carriers and high-pressure frozen in a Wohlwend Compact 3 (M. Wohlwend GmbH, Sennwald, Switzerland). Freeze substitution in acetone with 2% UA was initiated at −90°C and the temperature was increased 3°C/hour to −50°C. Then the cells were infiltrated and embedded in lowicryl HM20 resin polymerized by UV light. 50 nm thin sections were prepared and placed on formvar coated copper slot grids. Sections were stained with 2% uranyl acetate and Reynold’s lead citrate ^52^.

### Data acquisition

Cryo-EM/ET of human sperm was performed as described previously ^32^. Cryo-EM on bull sperm was performed on a Titan Krios operated at 300 kV with a Falcon3 detector (FEI, Eindhoven, Netherlands). Room temperature EM of bull spermatozoa thin sections was performed on a Tecnai T12 operated at 120 kV with a Ceta CMOS 16M camera (FEI, Eindhoven, Netherlands).

### Data analysis

Montages of all imaged cells were reconstructed by aligning overlapping cryo-electron micrographs. For the 23 cells whose tip was visible, the flagellum thickness was measured with IMOD ^53^ as the distance between the plasma membrane on each side of the flagellum. Three cells (out of 23) were excluded from the analysis because of large deformations of the flagellum plasma membrane (Supplementary Figure 2). Data points were chosen every 100 nm up to 5 µm away from the tip and every 1 µm up to 29 µm away from the tip. Thickness values of 18 to 20 sperm tails were included for each data point of the end piece and 5 to 18 tails were included for each data point of the principal piece, depending on what extent of the sperm tail could be imaged (Figure 2F). The distance between the flagellum tip and the end of the fibrous sheath could be measured in 19 cells. The average value was used to define the end piece length.

All cryo-electron tomograms were calculated and CTF-corrected with eTomo ^53^. sMTs and dMTs were independently counted in each tomogram (Supplementary Figure 1) and the extent of the singlet region was defined as the distance between the tip and the point where the number of sMTs equals that of dMTs. Due to the long singlet region of human spermatozoa, it is impossible to trace MTs back enough to distinguish between CP and extensions from dMTs, therefore in this work all MTs in the singlet region are collectively called sMTs.

## Acknowledgements

We thank Dr. J. Van Blerkom at Colorado Reproductive Endocrinology for providing human sperm samples from healthy patients. We thank Jonas Krantz at VikingGenetics for providing bovine sperm samples. We thank Lisa Larsson Berglund (Dept of Chemistry and Molecular Biology, University of Gothenburg) for helping with high-pressure freezing of bovine sperm samples. We thank Federica Tonolo (Dept of Biomedical Sciences, University of Padova) and Martin Palm (Dept of Chemistry and Molecular Biology, University of Gothenburg) for helpful comments on the manuscript. JHL was supported by a Sir Henry Wellcome Postdoctoral grant and a Swedish Research Council Young Investigator Grant (number 2015-05427). Electron microscopy of human samples was performed at the Boulder EM Services Core Facility in MCDB, University of Colorado, USA. Electron microscopy of bovine samples was performed at the SciLifeLab national Cryo-EM facility at Umeå Core facility for Electron Microscopy (UCEM), Umeå University, Sweden.

## Authors contributions

JLH designed and supervised the study. JTC and JLH prepared samples and all authors acquired data. DZ and JLH analyzed the data. DZ and JLH wrote the manuscript. All authors revised and approved the manuscript.

---

**Supplementary Figure 1. The number of singlet and doublet microtubules defines the singlet region.**

(A) The number of sMTs (blue data points) and the number of dMTs (orange data points) was counted along the flagellum. The striped background indicates the measured average extent of the singlet region and the grey background indicates the measured average extent of the end piece. (B) A complete axoneme has 2 sMTs and 9 dMTs.

**Supplementary Figure 2. Human sperm cells with morphological deformations.**

Montages of cryo-electron micrographs of three cells that were excluded from the analysis illustrated in Figure 2. The white arrows point at the morphological deformations. (A) The axoneme symmetry is disrupted in the principal piece. (B) Large bulge on the plasma membrane in the principal piece. (C) Bulge on the plasma membrane in the end piece.

**Supplementary video 1. Thickness of human sperm flagella.**

Visual representation of the gradual thickness changes at different points along the flagellum (from most distal to most proximal). Each frame represents one data point. For each data point the thickness was measured in at least 5 flagella. The diameter of each grey circle represents the measured thickness for one flagellum, the diameter of the black circle represents the mean thickness. The distance from the flagellar tip is specified for each frame. The red segments indicate the extent of the flagellar region in which each specific data point was acquired (end piece, singlet region; end piece, full axoneme; principal piece). Scale bar is 200 nm. Sperm cell cartoon not to scale.

**Supplementary video 2. Splitting of doublet microtubules into singlets in human sperm flagella.**

3D visualization of a dMT splitting into two sMTs. Each frame represents a 1 nm-thick tomographic view. Scale bar is 50 nm.

## Abbreviations

PCD: Primary Ciliary Dyskinesia
MT: Microtubule
sMT: Singlet Microtubule
dMT: Doublet Microtubule
CP: Central Pair
EM: Electron Microscopy
ET: Electron Tomography
IFT: Intraflagellar Transport

