## Supplementary figures and images for "Axonemal doublet microtubules can split into two complete singlets in human sperm flagellum tips"

### Supplementary Figure 1

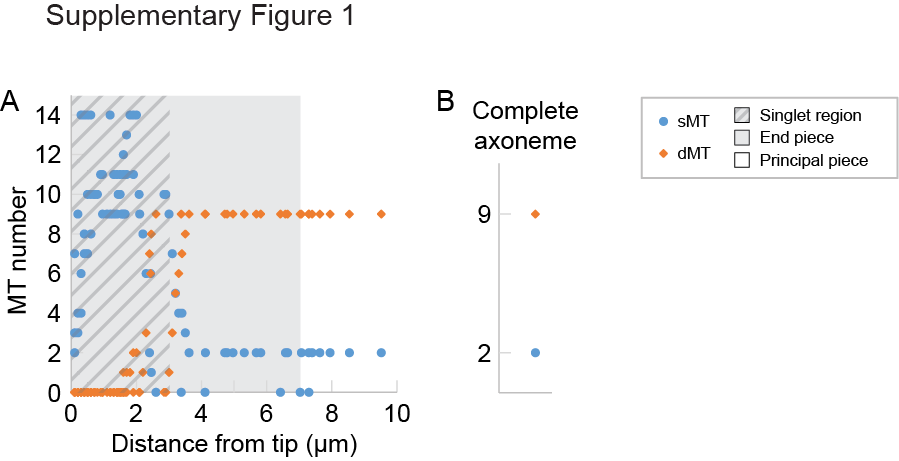

### Supplementary Figure 2

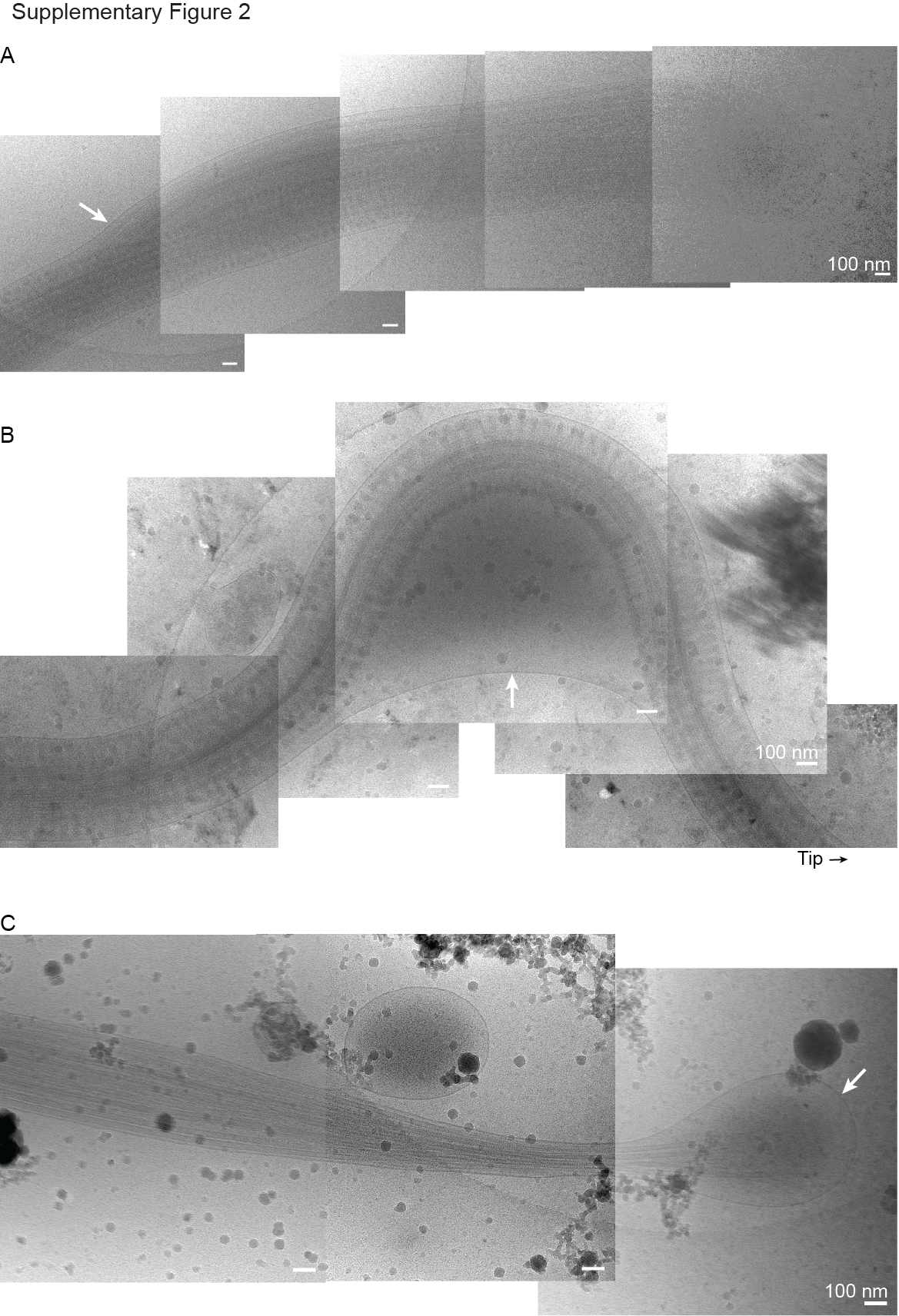
